# Inserting Pre-Analytical Chromatographic Priming Runs Significantly Improves Targeted Pathway Proteomics With Sample Multiplexing

**DOI:** 10.1101/2024.02.08.579551

**Authors:** Steven R. Shuken, Qing Yu, Steven P. Gygi

**Affiliations:** Department of Cell Biology, Harvard Medical School, 240 Longwood Ave, Boston, MA 02115, USA

## Abstract

GoDig, a recent platform for targeted pathway proteomics without the need for manual assay scheduling or synthetic standard peptides, is a relatively flexible and easy-to-use method that uses tandem mass tags (TMT) to increase sample throughput up to 18-fold relative to label-free targeted proteomics. Though the protein quantification success rate of GoDig is generally high, the peptide-level success rate is more limited, hampering the extension of GoDig to assays of harder-to-quantify proteins and site-specific phenomena. In order to guide the optimization of GoDig assays as well as improvements to the GoDig platform, we created GoDigViewer, a new stand-alone software that provides detailed visualizations of GoDig runs. GoDigViewer guided the implementation of “priming runs,” an acquisition mode with significantly higher success rates due to improved elution order calibration. In this mode, one or more chromatographic priming runs are automatically performed to determine accurate and precise target elution orders, followed by analytical runs which quantify targets. Using priming runs, peptide-level quantification success rates exceeded 97% for a list of 400 peptide targets and 95% for a list of 200 targets that are usually not quantified using untargeted mass spectrometry. We used priming runs to establish a quantitative assay of 125 macroautophagy proteins that had a >95% success rate and revealed differences in macroautophagy protein expression profiles across four human cell lines.

## INTRODUCTION

Proteins are the central enablers of the genome’s biological function, and their dysfunction and/or dysregulation plays a key role in nearly all diseases. In order to identify and characterize functional and dysfunctional biological states, quantification of proteins and their modifications is often required.^1^ One of the most powerful hypothesis-driven approaches to proteomic profiling is targeted mass spectrometry, in which specific peptide targets are separated using in-line HPLC and then detected and quantified via targeted monitoring, fragmentation, and mass analysis. Targeted proteomics is widely appreciated for its sensitivity, dynamic range, and quantitative accuracy and precision.^2,3^ Whereas traditional targeted proteomic methods such as parallel reaction monitoring (PRM) often need manual scheduling of MS2 scan acquisition for every target,^4,5,6^ we recently reported a novel targeted proteomic method, termed GoDig, which requires neither manual acquisition scheduling nor synthetic standard peptides and uses tandem mass tags (TMT) to increase sample throughput up to 18-fold.^1^ Using only a data-dependent acquisition (DDA)-based spectral and elution library and a target list, GoDig automatically monitors, identifies, and quantifies targets by measuring TMT reporter ion intensities. A given list of 200 proteins selected randomly from a publicly available library of >10,000 proteins can be targeted with GoDig with typical quantification success rates exceeding 80% in one injection (Figure S1). ^1^

Although the success rate of GoDig is generally suitable for biological pathway profiling (especially with multiple injections^1^), a more reliable single-injection success rate would be desirable. In addition, peptide-level success rates are often lacking, ranging from 50% to 60% (Figure S1), ^1^ limiting the utility of GoDig in targeting site-specific features. Here we describe two software engineering-based efforts to solve this problem: (1) the implementation of stand-alone software that visualizes successes and failures in GoDig acquisition and (2) the addition of a novel feature in GoDig that improves success rates by automatically improving elution order assignment accuracy.

GoDig determines in real time which targets are likely to be eluting by first identifying abundant “background” peptides in the sample using untargeted real-time search (RTS).^7^ Peptides identified by RTS are cross-referenced to the DDA-based library, which contains the relative binned retention times (elution order bins, EO bins) of the peptides in the library. In real time, EO bins of recently identified background peptides are aggregated into a “current calibrated EO bin.” When the current calibrated EO bin is within range of a given target’s EO bin, target monitoring is initiated. The size of the EO range is established by the user via an EO tolerance parameter, e.g., ±3 EO bins corresponding to (on average) ±1.5 min of real acquisition time in a 120-min run. The monitor scans initiate the process of identifying and quantifying the target. ^1^ When the current calibrated EO bin goes out of range, the target is no longer monitored.

Using new data visualization software (see below), we have found that when a target is not successfully quantified, one common explanation is that the target elutes outside the ±3-bin window due to chromatographic variability among systems. One solution is to widen the window; however, this decreases success rates by lowering duty cycle, especially with larger target lists. In light of this, we developed “priming runs,” which are runs solely dedicated to target identification and target EO bin correction that are collected before analytical runs and used to automatically correct EO bins during the experiment. Priming runs significantly improve peptide-level success rates in GoDig, making GoDig analysis readily applicable to single peptides and improving the reliability of pathway-spanning protein abundance assays.

## EXPERIMENTAL SECTION

### Spectral and elution library

DDA data for a spectral and elution library were acquired by analyzing the same 24 fractions of 4-cell-line peptides described in Ref. 1 with the same 120-min DDA method used in Ref. 1 with the only difference being the absence of a high-field asymmetric waveform ion mobility spectrometry (FAIMS) device. Spectra were also searched, and the peptide- and protein-level FDR controlled, using the same process.^1^ The resulting library contained 10,068 proteins and 145,623 unique peptides at 1% protein-level FDR and 0.08% peptide-level FDR.

### Four-cell-line TMTpro 16plex sample

The 4-cell-line TMTpro 16plex sample was prepared as follows. Human RPE1 (#CRL-4000), U2OS (#HTB-96), HCT116 (#CCL-247) and HEK293T (#CRL-3216) cells were purchased from American Type Culture Collection and grown in DMEM supplemented with 10% fetal bovine serum and 1% penicillin/streptomycin until 80% confluent. Cells were washed twice with ice cold PBS, pelleted, and stored at -80 °C until use. Cell pellets were lysed, and protein was precipitated, reduced, alkylated, digested, labeled with TMTpro reagents, combined, and cleaned as described previously.^8^

### General LC-MS/MS conditions

All liquid chromatography-tandem mass spectrometry (LC-MS/MS) experiments were carried out with an Oribitrap Eclipse tribrid mass spectrometer (ThermoFisher Scientific) equipped with an EASY-nLC 1200 system (ThermoFisher Scientific). All chromatography was conducted with a 30-inch column with a 100-μm inner diameter packed with ReproSil-Pur C18-coated beads 2.4 μm in diameter (Dr. Maisch GmbH, part no. r124.aq.0001) according to the FlashPack protocol.^9^ The column was heated to 60 °C and the total gradient flow rate was 550 nl/min. Peptides were separated using a 120-min gradient composed of buffer A (5% acetonitrile + 0.1% formic acid in water) and buffer B (95% acetonitrile + 0.1% formic acid in water): a 100-min linear gradient from 5% B to 25% B followed by a 10-min linear ramp to 41% B, then a 5-min ramp to 100% B followed by 5 min of isocratic flow at 100% B.

### Untargeted LC-MS/MS

Untargeted LC-MS/MS runs were conducted as follows. One μl of a 250 ng/μl suspension of 4-cell-line TMTpro 16plex was injected. The method length was 120 min. The solvent gradient described above was used. A real-time search-synchronous precursor selection-MS3 (RTS-SPS-MS3) method^7^ was constructed in XCalibur (ThermoFisher Scientific) with the following parameters and then used. MS1 scans were performed in the orbitrap at resolution = 120,000 with a scan range of 350–1500 Th. The quadrupole was used to isolate the scan range and the RF lens amplitude was set to 30%. The automatic maximum injection time feature was used, and the standard automatic gain control (AGC) target was 4x10^5^ charges. MS1, MS2, and MS3 spectra were all acquired in centroid mode with positive polarity.

Monoisotopic peaks were determined in peptide mode and an intensity threshold of 5x10^3^ charges was used for precursor selection. Ions of charge 2–5 were included in MS2 fragmentation. Dynamic exclusion duration was set to 90 min and began after 1 fragmentation event. The dynamic exclusion tolerance was ±10 ppm. All charge states and isotopes corresponding to the precursor were excluded. The entire scan cycle (including MS3) was limited to 3 s.

Selected precursors were isolated by the quadrupole with an isolation window 0.5 Th wide without offset. Collision-induced dissociation (CID) was performed in the ion trap with a fixed normalized collision energy of 35% for 10 ms and activation Q parameter of 0.25. MS2 scans were acquired in the ion trap with Turbo scan rate and automatic scan range. The standard AGC target of 1x10^4^ charges was used with a maximum injection time of 35 ms.

Real-time search (RTS)^7^ was executed in XCalibur (ThermoFisher Scientific). RTS was performed on MS2 scans using a concatenated forward and reverse human reference proteome based on the FASTA file available from Uniprot (updated December 21, 2018). Tryptic peptides excluding KP/RP sites were included, with cysteine carbamidomethylation and lysine and peptide N-terminus TMTpro modifications as static modifications and methionine oxidation as a variable modification. A maximum of two missed cleavages and two variable modifications were considered. Protein close-out was activated after identifying two peptides per protein. The maximum search time was 100 ms. The Xcorr threshold was 1.4, the dCn threshold was 0.1, and the precursor ppm tolerance was 20 ppm for z = 2 and 10 ppm for z = 3 or 4. Peptides matching reverse and contaminant proteins were excluded.

Only fragment ions with m/z between 400 and 2000 Th were included in SPS. Values of m/z from 50 Th below to 5 Th above the precursor mass were excluded. Fragment ions with TMTpro tag loss were excluded. Up to 10 SPS ions were selected. Peptides were isolated within a 1.2-Th window and each fragment was isolated with SPS within a 2-Th window without offset. Fragments were subjected to HCD with a normalized collision energy of 55%. Reporter ions were analyzed in the orbitrap at a resolution of 50,000. A scan range of 110 to 1000 Th was used. A custom AGC target of 300% was used with a maximum injection time of 200 ms.

### GoDig LC-MS/MS analysis

Unless otherwise noted, GoDig runs were performed as follows. One μl of a 250 ng/μl suspension of 4-cell-line TMTpro 16plex was injected. The method length was 120 min. The solvent gradient described above was used. The 4-cell-line-based library described above was used. For RTS, the human reference proteome from Uniprot (updated December 12, 2021) was concatenated with reversed versions of the proteins and indexed in the GoDig software using the TMTpro modification.^1^ Lysine and peptide N-terminal TMTpro labeling and cysteine carbamidomethylation were included as static modifications and methionine oxidation was included as a dynamic modification. Elution order (EO) calibration was performed every 15 s and the top 6 MS1 features were included in EO calibration MS2. The method length was set to 120 min. Peptides were monitored using ion trap CID-MS2 scans with an RF lens parameter of 30%, quadrupole isolation width 0.5 Th, normalized collision energy 34.0%, AGC target of 10,000 charges, maximum injection time 120 ms, and scan rate set to Normal. The monitor MS2 fragment peak match tolerance was ±0.15 Th; monitor MS2 scans with ≥6 matched fragment peaks triggered identification MS2 (IDMS2). For identification, precursors were isolated with the quadrupole using a 0.5-Th window and then subjected to CID using a normalized collision energy of 34.1%. The IDMS2 AGC target was 1x10^5^ charges and the maximum injection time was 600 ms. Fragments were analyzed in the orbitrap at a resolution of 15,000. The fragment match tolerance was 15 ppm and a minimum of 4 fragment peaks was required for a cosine score comparison. The cosine match score threshold was 0.8. For MS3, the same filters as for untargeted SPS (see above) were used for SPS ion selection (MS2 ions 50 ppm below or 5 ppm above precursor m/z or outside the 400–2000-Th range were excluded). Up to 4 SPS ions were isolated with an isolation window width of 0.8 Th. Isolated fragments were subjected to HCD at a normalized collision energy of 45% and MS3 scans were acquired in the orbitrap at a resolution setting of 50,000. An MS3 prescan was used to calculate the injection time. The MS3 AGC target was 250,000 charges and the maximum injection time was 1000 ms.

### Software Development

GoDig was developed in C# using the instrument API version 2.0 (June 9, 2020) (ThermoFisher Scientific) and .NET Framework 4.7.2 with Windows Forms. GoDigViewer was developed in C# using the RawFileReader NuGet package version 5.0.0.7 (ThermoFisher Scientific) and .NET Core 5.0 with Windows Presentation Foundation.

## RESULTS

### GoDigViewer

We developed a new software for the visualization of GoDig runs called GoDigViewer (Figure 1A). In GoDigViewer, all MS2 and MS3 events in a run can be simultaneously visualized, giving a holistic view of the experiment (Figure S2); counts and success rates at the levels of scans, charge state-specific precursors, peptides, and genes are reported (Figure S3). In addition, a particular target precursor can be selected and analyzed in a separate view, where all spectra relevant to the target may be visualized and a fragment ion chromatogram can be inspected (Figure S4). Visualization of real-time EO calibration is also provided, at both global and target-specific levels (Figure S5–S6).

**Figure 1.**
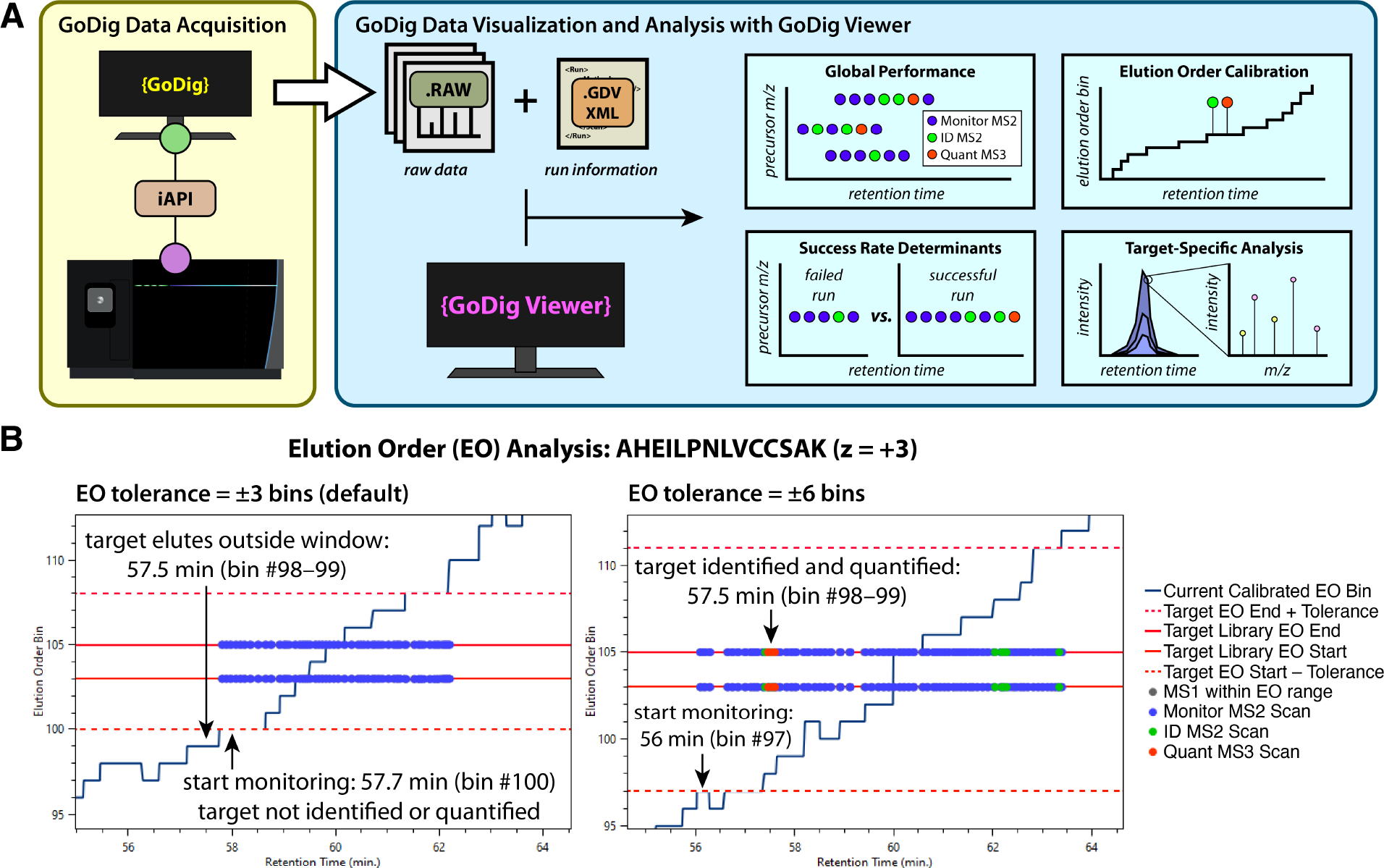
GoDigViewer reveals out-of-bounds elution as a GoDig success rate determinant. **A**. GoDig and GoDigViewer workflow. During a GoDig LC-MS/MS run, two files are generated: a GoDigViewer-compatible XML file (file extension .gdvxml) is generated in addition to the raw file. GoDigViewer imports these two files and provides a suite of visualization features (for exemplary screenshots see Figures S2–S6). **B**. Annotated GoDigViewer screenshots showing an example of out-of-bounds elution causing a target to be neither identified nor quantified (left) and this issue being fixed by alteration of the elution order (EO) tolerance parameter (right). The blue trace illustrates the current calibrated EO bin established by real-time search (RTS) and cross-referencing RTS results to the elution library. Gray dots (mostly obscured) represent within-bounds MS1 scans; blue dots represent monitor scans; green dots represent attempted identification MS2 (IDMS2) scans triggered by monitor scans; red-orange dots represent MS3 scans triggered by successful IDMS2 scans. The central two horizontal red lines represent the range of EO bin values of the target in the library, and the other two represent the window established by the EO tolerance. In this example, the target was successfully identified 5 bins away from the expected elution point, suggesting that a tolerance of ±3 EO bins was insufficient for this target.

### GoDigViewer illuminates out-of-bounds elution as a source of failure in GoDig

GoDigViewer can be used to straightforwardly diagnose and fix GoDig failures using an iterative process: after each run, a new run can be performed with altered parameters in an attempt to detect and quantify missed targets. For example, if targets elute outside of the default EO windows, the windows can be widened to detect these targets (Figure 1B). This phenomenon, which we call “out-of-bounds elution,” was repeatedly observed during the implementation and testing of GoDigViewer. To test the efficacy of widening EO windows in solving this problem, we analyzed a TMTpro-labeled sample containing peptides from 4 different cell lines (Figure 2A) with different EO tolerances. We curated a target list by analyzing fifteen identical untargeted real-time search-synchronous precursor selection-MS3 (RTS-SPS-MS3) runs, each performed on 250 ng of the 4-cell-line TMTpro 16plex sample and selecting 400 of the most commonly detected and quantified precursors in the dataset (“super fliers”) (Figure 2B). Since these precursors generate reproducible signals and fragment well, it is presumable that a failure to detect one of these targets with GoDig would likely result from out-of-bounds elution. Although widening windows can identify some out-of-bounds-eluting targets, these benefits are ultimately diminishing, because acquisition becomes more crowded (lower duty cycle) as EO tolerances are increased (Figure S7).

**Figure 2.**
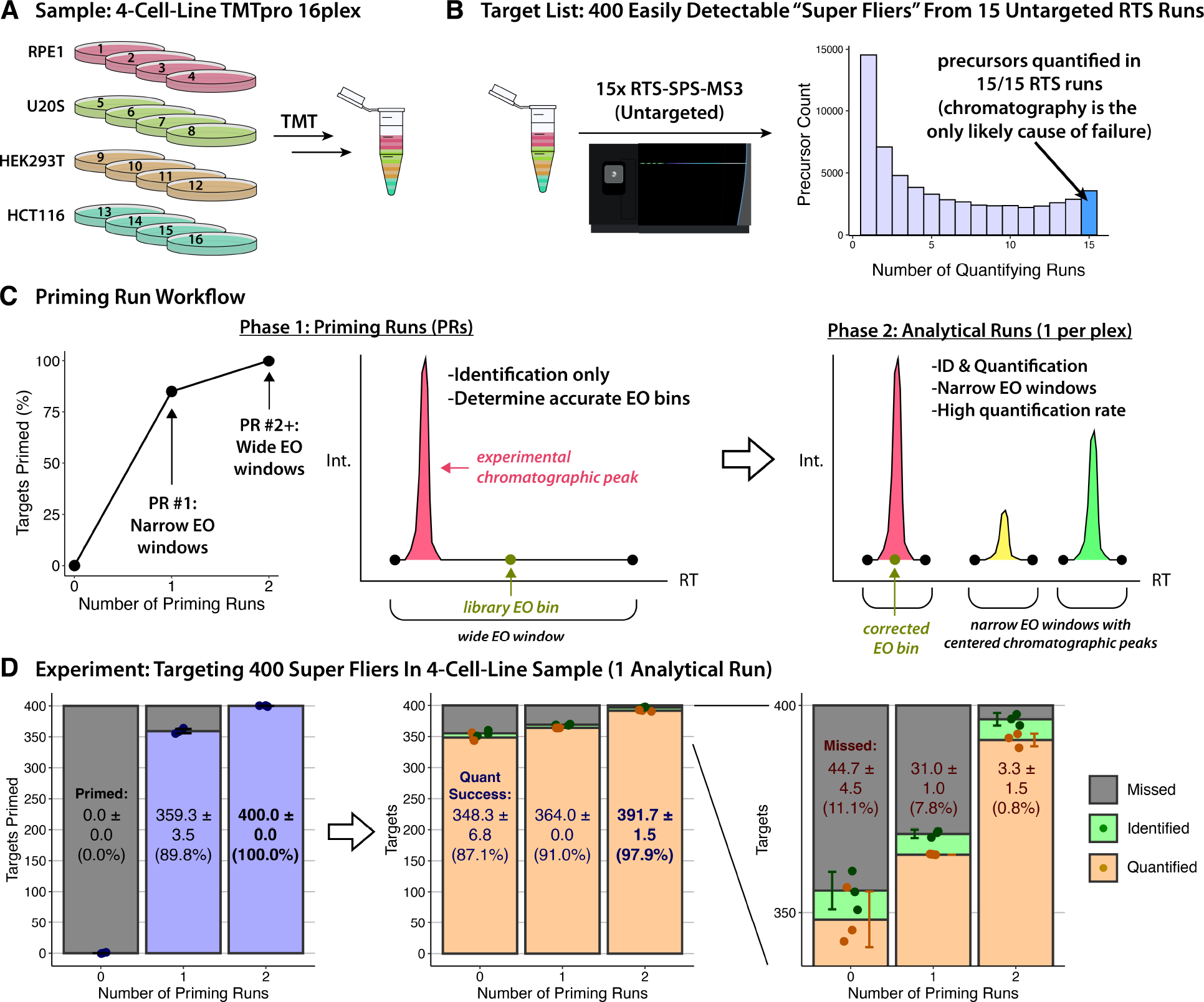
Priming runs. **A**. Four-cell-line TMTpro 16plex sample used for priming run implementation, testing, and evaluation. **B**. Selection of “super flier” targets for which out-of-bounds elution is a major likely cause of failure to quantify. **C**. Priming run concept. First, identification-only GoDig runs with widening EO tolerances (windows) are performed in order to determine an empirical EO bin which is then used as the corrected EO bin for the analytical runs. “Primed” means identified and corrected. **D**. GoDig analysis of the 4-cell-line sample targeting 400 easily detected precursors with and without priming runs. “Identified” means the cosine match threshold and spectral quality filters were passed and MS3 was triggered. “Quantified” means the summed TMT signal-to-noise ratio exceeded 160 and isolation purity of SPS ions exceeded 50%. An analytical run was performed after a set number of priming runs (i.e., 0, 1, 2). Peptides identified (primed) during the priming run(s) are plotted in the left panel and peptides identified and quantified during the analytical run are plotted in the middle and right panels.

### Priming runs raise GoDig quantification success rates to >97%

To robustly address the issue of out-of-bounds target elution, we conceived priming runs (Figure 2C). In a priming run, peptides are identified (not quantified, i.e., MS3 scans are not executed) and an empirically corrected EO bin is determined for each identified target. For the first priming run, the default analytical EO tolerance of ±3 bins is used, whereas in the second (and later) priming run(s), a wider EO tolerance, e.g. ±15 bins (±7.5 min on average), is used. If there are multiple successful IDMS2 scans for a target, the median current calibrated EO bin among them is used as the corrected EO bin for that target in subsequent analytical run(s). The corrected EO bins are automatically used in the analytical run(s) and are written to a file for use in later GoDig experiments.

In principle, a set of priming runs can identify all of the targets in a list of detectable precursors. This is accomplished by “run-wise close-out” which is automatically enabled during priming runs: once a target is identified, it is excluded from subsequent priming run(s). Thus, each priming run targets precursors that were missed by the previous priming run(s), enabling an exhaustive analysis. Indeed, in 2 priming runs, all 400 super fliers were reproducibly successfully “primed” (detected and empirical EO bins established) in 2-hr analyses of 250 ng of the 4-cell-line sample (Figure 2D). Inserting priming runs before analytical runs increased the average quantification success rate from 87.1% to 97.9% (Figure 2D). Importantly, nothing was changed in the analytical run parameters; priming runs were merely conducted beforehand automatically by the GoDig software. When using priming runs, quantification is consistent with GoDig without priming runs (Figure S8–S9). To show that the performance boost is sustained over multiple analytical runs, as would be the case for GoDig experiments with more than one TMT plex, we performed a GoDig experiment targeting these 400 targets with 2 priming runs and 4 consecutive analytical runs. Again, all 400 targets were primed, and the quantification success rate, averaging 97.9%, was stable across the 4 analytical runs (Figure S10).

### Priming runs increase elution order bin accuracy and precision

Using GoDigViewer, we investigated how these improved success rates are achieved. An example of the effect of priming runs on a particular target is shown in Figure 3. In this example, performing GoDig without priming runs missed the target by out-of-bounds elution; the second priming run, with an EO tolerance of ±15 bins (corresponding to ±7.5 min on average), identified the target and corrected the EO bin, enabling the target to be successfully identified and quantified in the subsequent analytical run.

**Figure 3.**
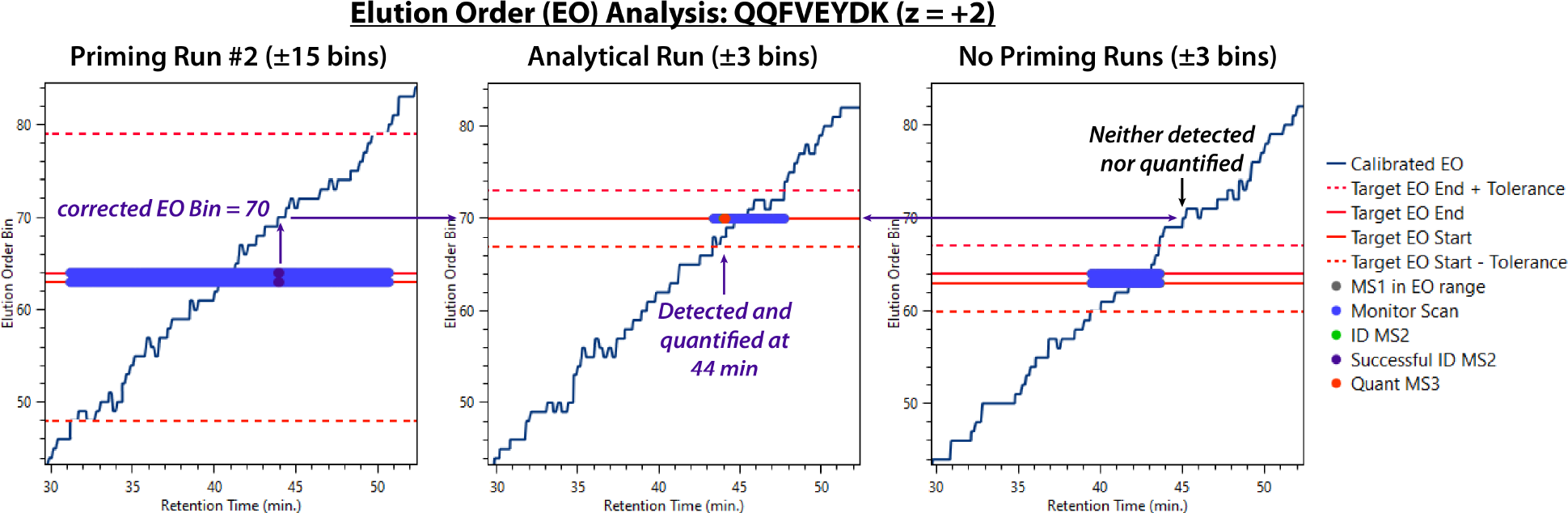
Example effect of priming runs on a target. Plots were generated in GoDigViewer with the same data from Figure 2D. The dark purple dot represents the successful IDMS2 during the priming run. In this example, a target with a library EO bin range of 63–64 was detected during the 2^nd^ priming run at current calibrated EO bin = 70. The target was successfully detected and quantified in the subsequent analytical run (middle panel), whereas, without priming runs the EO window spanned bins 60–67, missing the target.

A global visualization in GoDigViewer of the same runs shows that overall, priming runs result in more accurate and precise EO bins (Figures 4, S11, and S12). Whereas Figure 4 plots the full library target EO ranges, de-emphasizing the differences between priming runs #1 and #2, Figures S11–S12 only plot the minimum distance from the current calibrated EO bin and the target EO range, showing the necessity of the wide EO tolerance in priming run #2. Figure S12 illustrates the increased accuracy of corrected EO bins by plotting these minimal EO bin errors.

**Figure 4.**
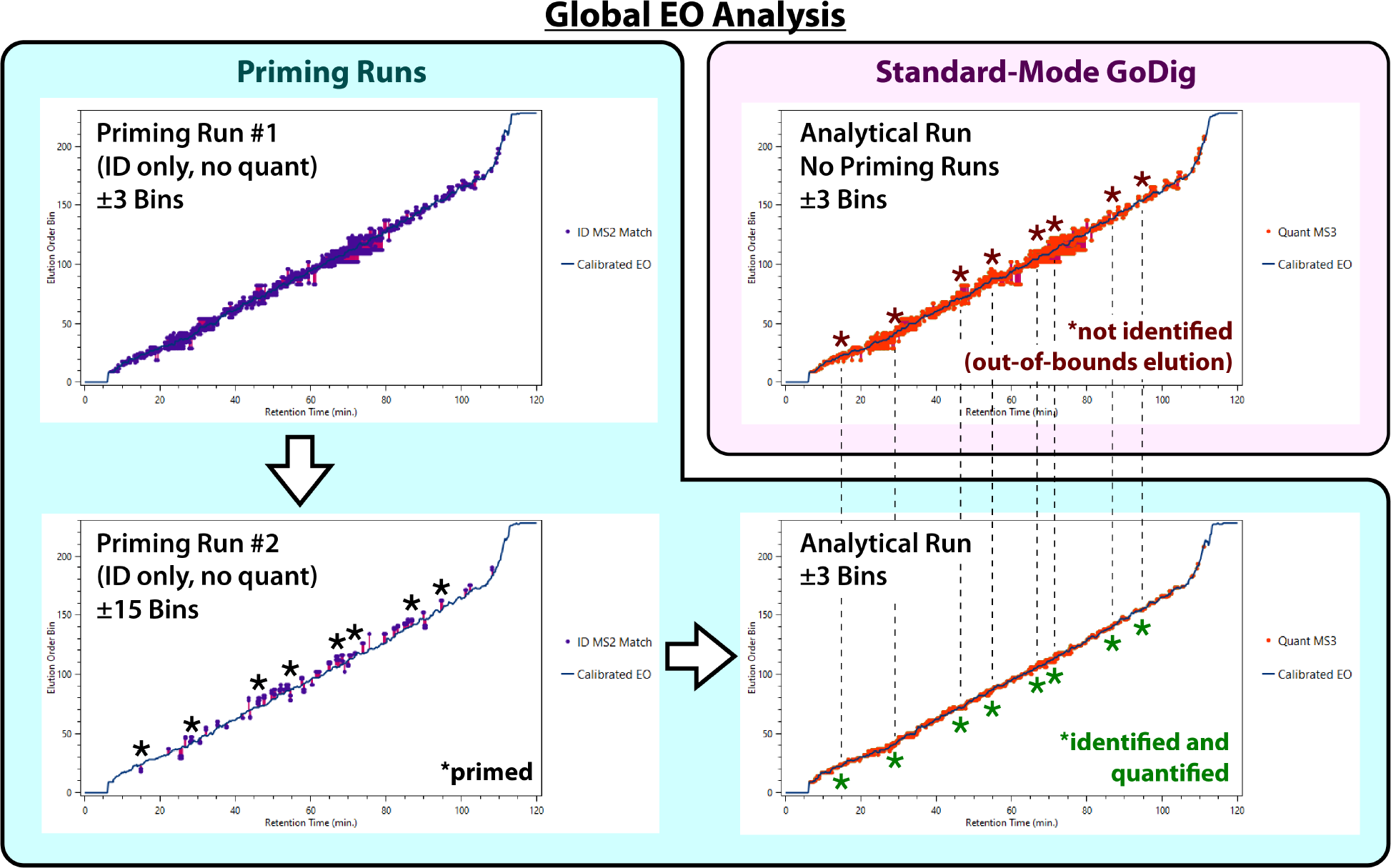
GoDigViewer screenshots illustrating the global effect of priming runs on EO bin accuracy and precision. Purple dots represent successful identification events in priming runs whereas and red dots represent quantification events in analytical runs which are triggered by successful identification events. Vertical magenta lines show the EO bin error. Without priming runs, these EO errors are relatively large (top-right); during priming runs, they are similar (top-left) and then intentionally allowed to be larger (bottom left); in an analytical run preceded by priming runs, EO bin errors are greatly reduced (bottom right). Some examples of targets that were not identified without priming runs because of out-of-bounds elution, primed during priming run #2, and subsequently quantified are indicated with asterisks and dotted lines.

To test the idea that priming runs enable an analytical tolerance below ±3 bins, we repeated the 2-priming-run GoDig experiment depicted in Figure 2D in triplicate and then used the resulting corrected EO bins to perform the same GoDig analysis with an EO tolerance of ±2 bins and ±1 bin (±1 min and ±30 s, respectively). We observed a slight decrease in performance from 97.8% with ±3 bins 96.5% with ±2 bins, whereas with ±1 bin we quantified 86.2% (Figure S13). Since 96.5% is a useful success rate for most studies, this approach may be helpful for studies targeting larger lists of peptides in which increased time efficiency might have a stronger effect on the success rate.

### Priming run-based quantification rates exceed 95% for peptides quantified in less than half of untargeted runs

Having demonstrated the performance increase that comes with priming runs, we tested the priming run feature on a more difficult set of targets: targets that were quantified in less than half of the fifteen untargeted RTS-SPS-MS3 runs described above. We curated a list of 200 such precursors and targeted them with GoDig using priming runs (Figure 5). Whereas a single RTS-SPS-MS3 run would on average quantify only 46.7% of these precursors, GoDig without priming runs averaged 78.8% quantification success, and with 2 priming runs, the success rate averaged 95.4% over 4 runs. Thus, priming runs enable reliable detection and quantification at the peptide level, even with peptides that are not usually quantified with untargeted mass spectrometry.

**Figure 5.**
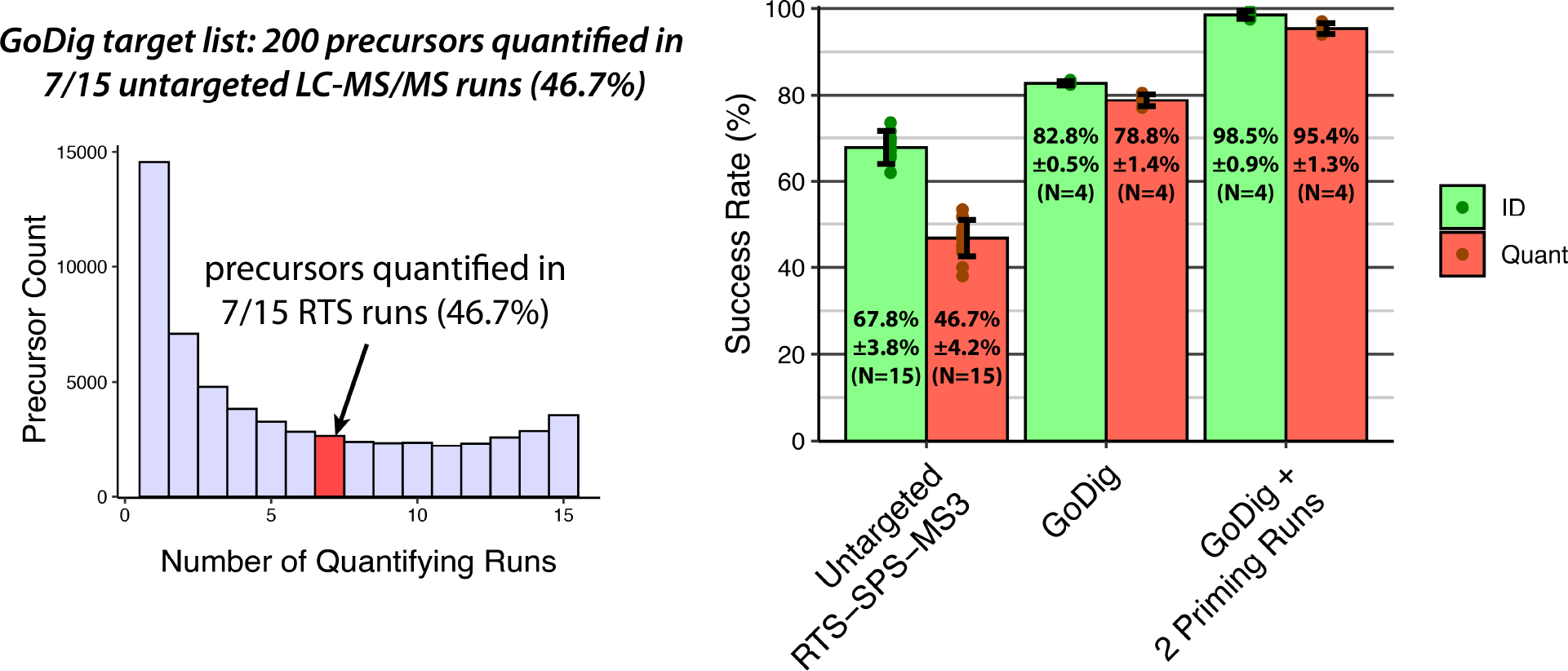
Using priming runs to target precursors that were identified in less than half of untargeted RTS-SPS-MS3 runs. A target list of 200 precursors was curated wherein all of the precursors were quantified in 7 of 15 untargeted runs. Numbers reported for each condition are, in order, the mean success rate, standard deviation, and number of replicate runs. Green data represent identification (successful IDMS2) and red-orange data represent quantification (summed TMT signal-to-noise ratio exceeds 160).

### A priming run-based macroautophagy pathway assay reveals cell line-specific differences with a >95% success rate

To showcase the utility of priming runs in studying biologically relevant targets, we used priming runs in “targeted pathway proteomics,” a technique which quantifies a set of proteins representing a biological pathway.^1^ Autophagy, defined as any cellular degradative pathway that involves the delivery of cytoplasmic cargo to the lysosome,^10^ is a pathway with special relevance to homeostasis, aging, and disease.^11^ At least three types of autophagy are known: chaperone-mediated autophagy, microautophagy, and macroautophagy, in which a double-membrane-bearing phagophore engulfs cargo, closes to form an autophagosome, and then fuses with endosomes or lysosomes for degradation (Figure 6A).^12^ Macroautophagy plays a role in the degradation of α-synuclein in Parkinson’s disease, huntingtin in Huntington’s disease, and hyperphosphorylated tau and amyloid beta-42 in Alzheimer’s disease, and activation of macroautophagy via TORC1 inhibition has shown substantial benefits in various animal models of these diseases and more.^13,14^ Using the QuickGO web tool (https://www.ebi.ac.uk/QuickGO/term/GO:0016236), we found in the human genome 245 genes with the gene ontology (GO) term for macroautophagy (GO:0016236). Our 4-cell-line-based GoDig library contains at least two unique peptides for 161 of these genes. For each of these 161 genes, we selected two peptides to target, primarily based on how many times they were quantified in the 15 untargeted RTS runs and secondarily based on MS2 spectral quality in the library (using comparable statistical scores based on linear discriminators).^3,15^ Over the course of four priming runs, 125 of these genes were primed by detection of both precursors (Figure S14). Using the corrected EO bins from these priming runs, we targeted these 125 genes in the 4-cell-line mixture for four analytical runs, identifying 98.8% ± 0.5% and quantifying 95.6% ± 1.0% compared to 87.0% ± 1.2% with uncorrected EO bins and 35.7% ± 2.3% in untargeted RTS (Figure 6B).

**Figure 6.**
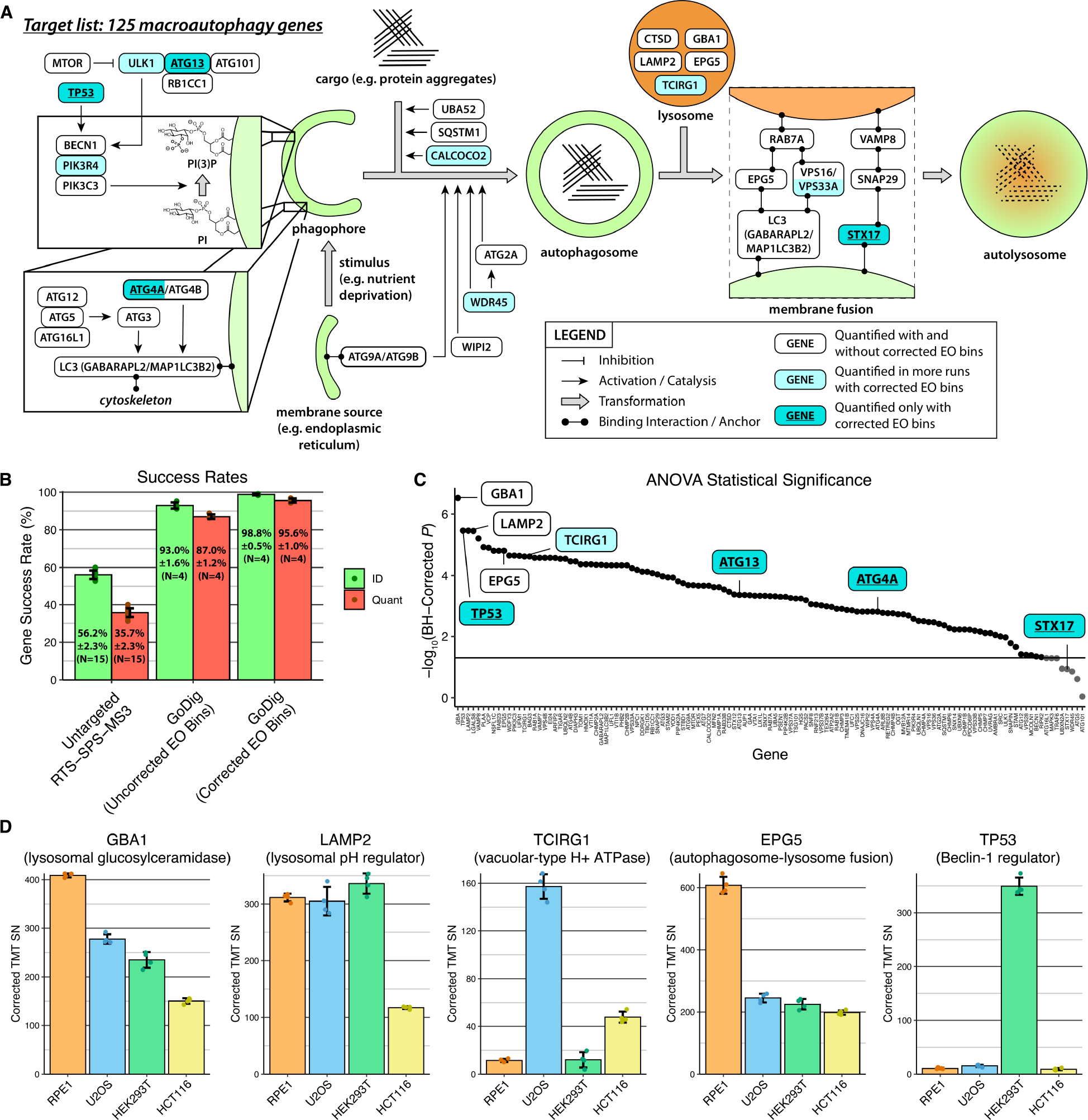
A priming run-based macroautophagy GoDig assay. **A**. Simplified diagram of macroautophagy pathway with a selection of genes shown. All genes shown are in the 125-gene assay and were quantified by GoDig runs using EO bins corrected by priming runs. Each gene appears exactly once except LC (GABARAPL2/MAP1LC3B2) which appears twice. **B**. Same as Figure 5 except that success rates are calculated at the gene level and the target list is 125 macroautophagy genes. **C**. Negative logarithm of Benjamini-Hochberg (BH)-corrected ANOVA *p*-value for each gene. Genes highlighted in **A** and **D** are emphasized. **D**. Quantitative data for selected genes are shown as corrected TMT reporter ion signal-to-noise (SN) ratios.

The corrected EO bins enabled quantification of 8 genes which were never quantified without priming runs, four of which are highlighted in Figure 6A: ATG13, a member of the ULK complex; TP53, a regulator of the central protein Beclin-1; ATG4A, which cleaves the C-terminus of microtubule-associated protein light chain 3 (LC3) proteins, an essential preprocessing step; and STX17, which plays a role in lysosome-autophagosome membrane fusion.^16,17^ In addition, 21 genes were quantified in more runs with corrected EO bins than without (including 10 highlighted in Figure 6A) and 112 genes were quantified in all 4 runs with corrected bins compared to 99 with uncorrected bins.

The highly complete dataset produced by GoDig with corrected EO bins contains protein-level gene expression data across the 4 cell lines for these 125 genes. To get a global sense of variation in macroautophagy protein expression across these cell lines, we performed one-way ANOVA tests, most of which yielded significant *p*-values after Benjamini-Hochberg correction, including some proteins not quantified with uncorrected EO bins (Figure 6C).^18^ The data for a selection of the most significant genes are plotted in Figure 6D. The quantitative data agree with results obtained previously using untargeted RTS-SPS-MS3 analysis of a 4-cell-line mixture fractionated with offline HPLC (Figure S15).^1^ Three lysosomal genes, GBA1, LAMP2, and TCIRG1, had significantly different cell line-specific profiles, supporting claims that cells from different lines maintain different lysosomal proteomes.^19^ TCIRG, a subunit of a vacuolar-type H+-transporting ATPase, is known to be important in osteoclast generation in bone;^20^ this may explain why its expression is more than 3x greater in the osteoblast-derived U2OS line. EPG5 facilitates the necessary STX17-VAMP8 interaction during autophagosome-lysosome fusion; patients with EPG5 mutations develop Vici syndrome, characterized by severe developmental dysfunction in the brain, heart, and eyes.^21,22^ The importance of EPG5 in ocular development and function may explain its increased expression in the retina-derived RPE1 cells. In addition to its role in regulating autophagy, TP53, also known as p53, is a well-known tumor suppressor and is mutated in most human cancers;^23,24^ because the HEK293T line, unlike the others, is not a cancer line, we measured much higher levels of p53 in HEK293T cells. These data confirm that with priming runs, GoDig can precisely and accurately measure protein abundance differences not measurable with either untargeted data-dependent acquisition or priming run-free GoDig.

## DISCUSSION

In targeted proteomics, the success rate of an assay must be high in order to be useful in large-scale biological studies. In the case of protein quantification, standard-mode GoDig is suitable for many studies; however, in this work, we have described an acquisition mode that is not only superior for protein quantification but also applicable to the quantification of individual peptides. This opens up a large potential space of GoDig studies, including studies of phosphorylation-based signaling pathways,^25^ proteolytic cleavage sites such as those induced by limited proteolysis-mass spectrometry,^26^ or sites of covalent drug binding such as reactive cysteine residues.^27^

We arrived at the concept of priming runs using a new software that enables detailed visualization of the of the GoDig raw data acquisition process. Indeed, most software packages that analyze any type of mass spectrometry data do not reveal real-time decision making or provide failure analyses with the level of detail exhibited by GoDigViewer. Using GoDigViewer, additional features will be implemented to further increase success rates for larger and more difficult target sets.

We used priming runs not only to increase success rates but also to develop a quantitative assay of 125 macroautophagy proteins. This process exemplified the ease of use of GoDig: once the automatic priming runs were complete, the assay was ready for use, and it can be used in future studies on systems where macroautophagy is particularly relevant, such as models of aging or neurodegeneration. In the future, priming runs will be used in the development of additional high-value assays for particular pathways of interest. Such assays will enable routine investigation of critical biological and pathological processes with unprecedented sensitivity, reliability, and sample throughput.

## ASSOCIATED CONTENT

### Supporting Information

Supplementary figures S1–S15. AUTHOR INFORMATION

### Author Contributions

Q.Y. created the GoDig platform and advised on the project. S.R.S. and S.P.G. conceived the project. Q.Y. made the 4-cell-line TMTpro 16plex sample. S.R.S. implemented GoDigViewer, implemented the priming run feature in GoDig, performed untargeted and targeted mass spectrometry experiments, analyzed and plotted the data, and wrote the manuscript. S.R.S., Q.Y., and S.P.G. edited the manuscript.

### Funding Sources

NIH grant GM067945 to SPG.

## ACKNOWLEDGMENT

The authors thank Jesse Canterbury and William Barshop (ThermoFisher Scientific) for help with the instrument API, João Paulo for all-around advice with instrument issues, and members of the Gygi Laboratory for helpful conversations and support.

