## Supplemental Data 1 for "Inserting Pre-Analytical Chromatographic Priming Runs Significantly Improves Targeted Pathway Proteomics With Sample Multiplexing"

### Supplementary Figures

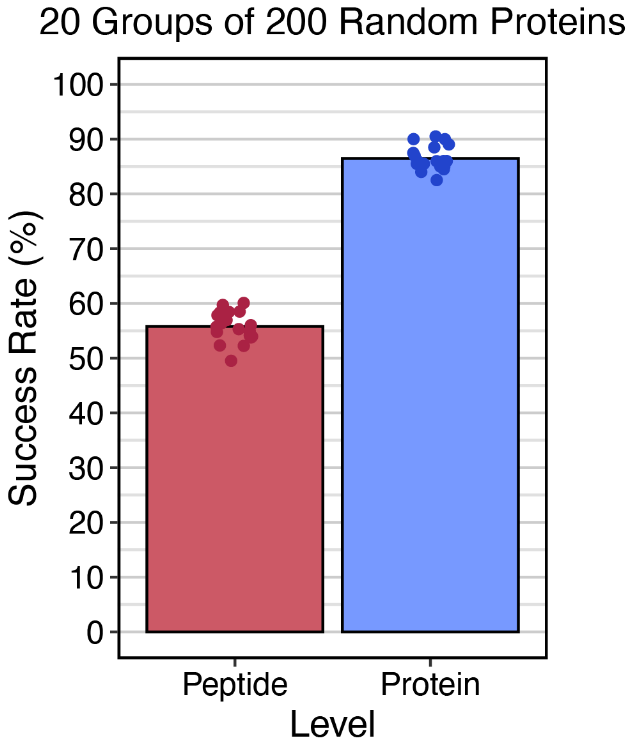


**Figure S1. Reanalysis of GoDig data from Yu et al. 2023 (Ref. 1).** The success rate is defined as the number of targets with summed TMT reporter ion signal-to-noise ratios greater than 10 per channel divided by the number of targets in the list.

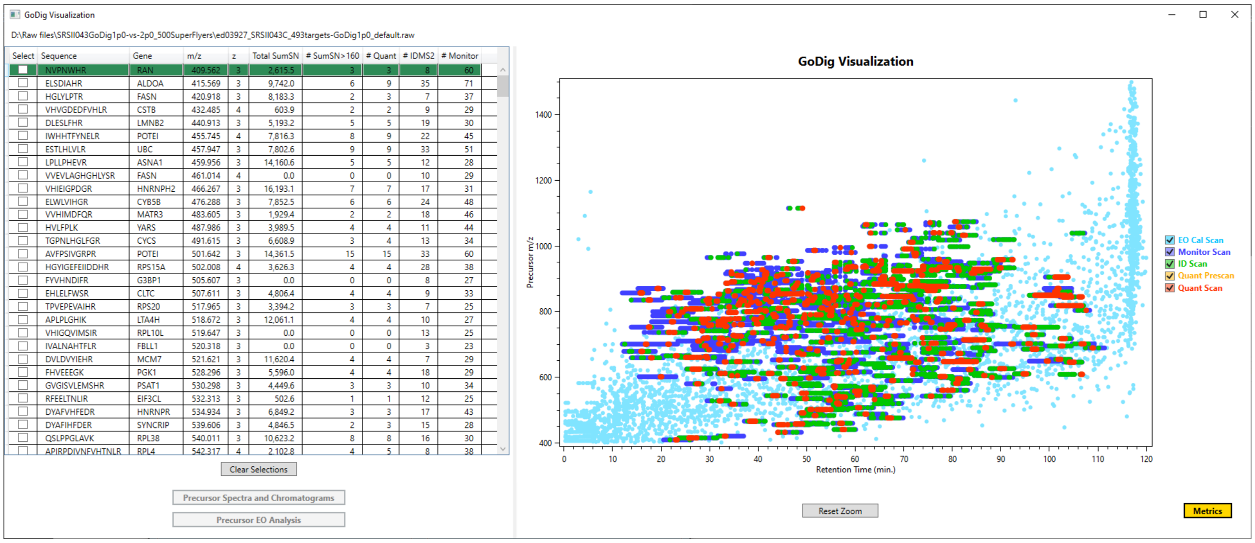


**Figure S2. Run visualization in GoDig Viewer.** The left panel is a grid listing all targets along with their gene symbols, m/z values, charge states, total accumulated TMT reporter ion peak signal-to-noise sums (SumSN values), number of MS3 scans with SumSN exceeding 160, number of MS3 scans ("# Quant"), number of ID MS2 scans, and number of monitor scans. The right panel is a plot of all MS2 and MS3 scans in the run with the precursor m/z on the y-axis. EO Cal Scan = elution order calibration MS2 scan; Monitor Scan = monitor scan (ion trap MS2 or MSX-SIM scan); ID Scan = identification orbitrap MS2 scan; Quant Prescan = automatic gain control prescan for MS3; Quant Scan = SPS-MS3 scan.

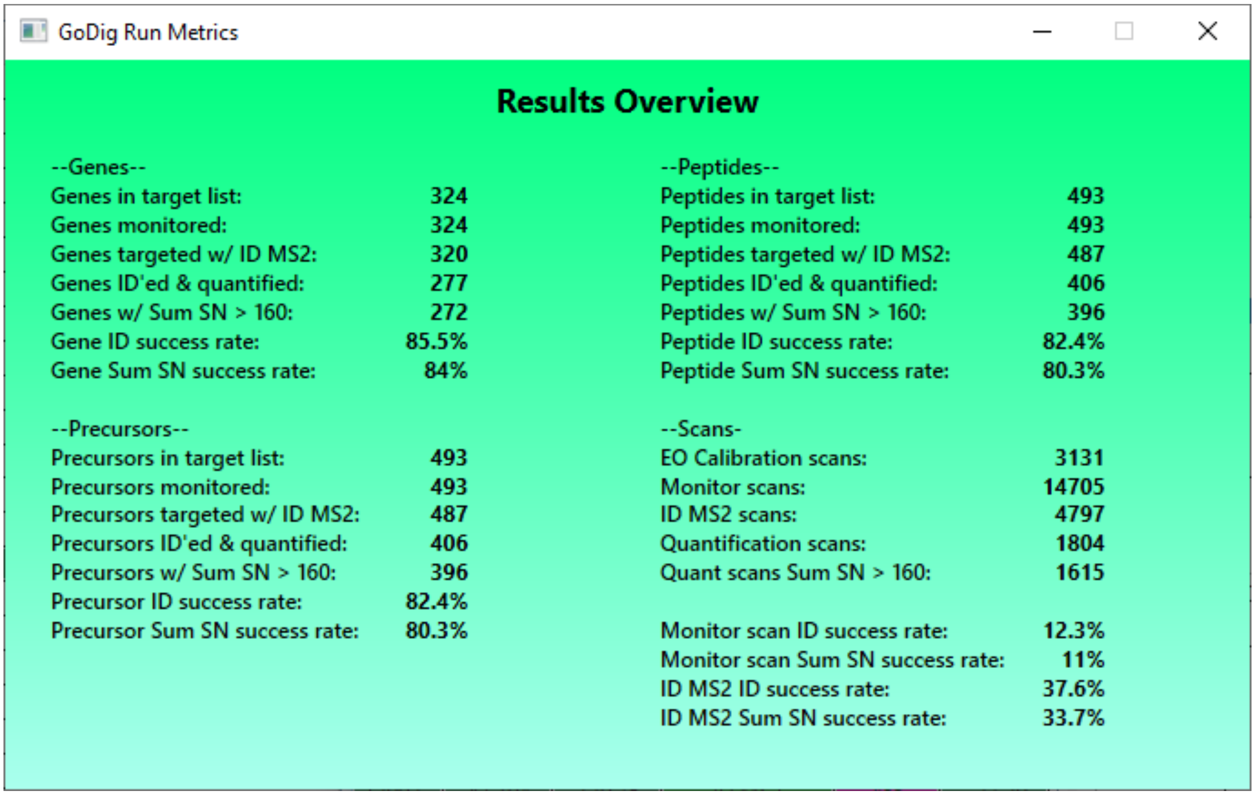


**Figure S3. Example of the Metrics pane in GoDig Viewer.** "ID'ed & quantified" signifies that an ID MS2 spectrum matched the library spectrum and triggered an MS3 scan. "Precursor" signifies a charge state-specific peptide ion. "Sum SN" refers to the total sum of TMT reporter ion signal-to-noise ratios (summed across multiple MS3 scans if available).

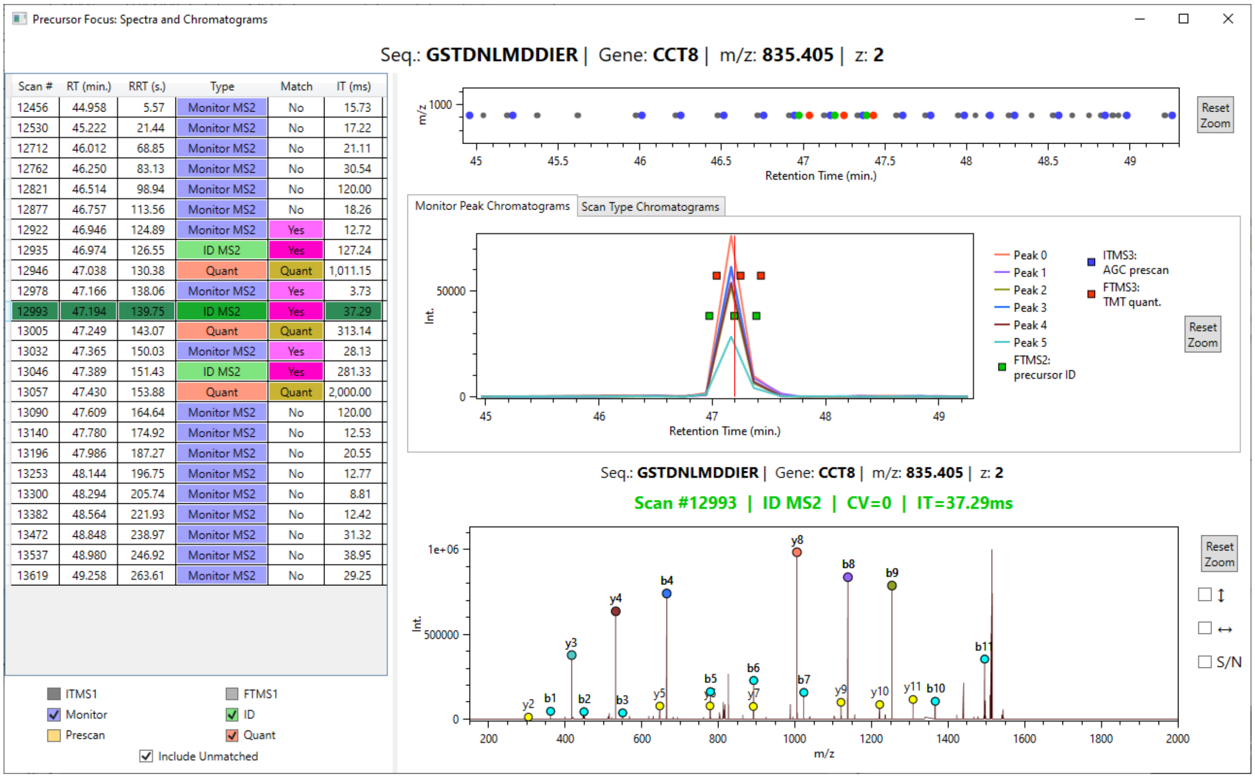


**Figure S4. The Precursor Focus pane in GoDig Viewer.** Left panel: list of all scans relevant to the target, which can be filtered by scan type using the check boxes at the bottom left. RRT = relative retention time (RT relative to the beginning of target monitoring). IT = injection time. Right-top panel: same as the plot in Figure S2 except only scans pertaining to this target are included and MS1 scans are added as gray dots. Right-center panel: chromatogram consisting of major fragment peak intensities from monitor MS2 scans. Green squares are successful ID MS2s and red-orange squares are MS3s. Right-bottom panel: spectrum of scan selected from list. CV = FAIMS compensation voltage.

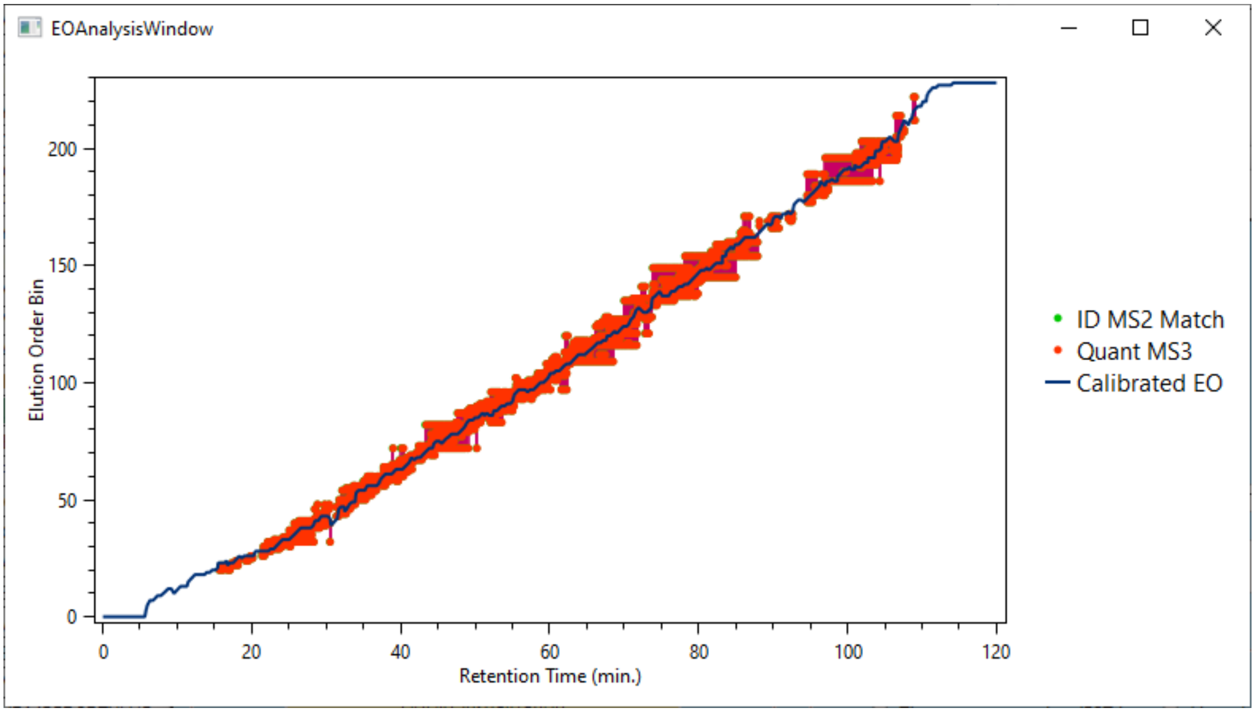


**Figure S5. The Global EO Analysis pane in GoDig Viewer.** The blue line plot illustrates the current calibrated EO bin established by real-time search (RTS). Green dots (mostly obscured by the corresponding red-orange dots) represent successful IDMS2 scans and red-orange dots represent the subsequent MS3 scans. Each dot is attached to the blue curve by a vertical purple line that illustrates the difference between the current calibrated EO bin and the target's library EO bin.

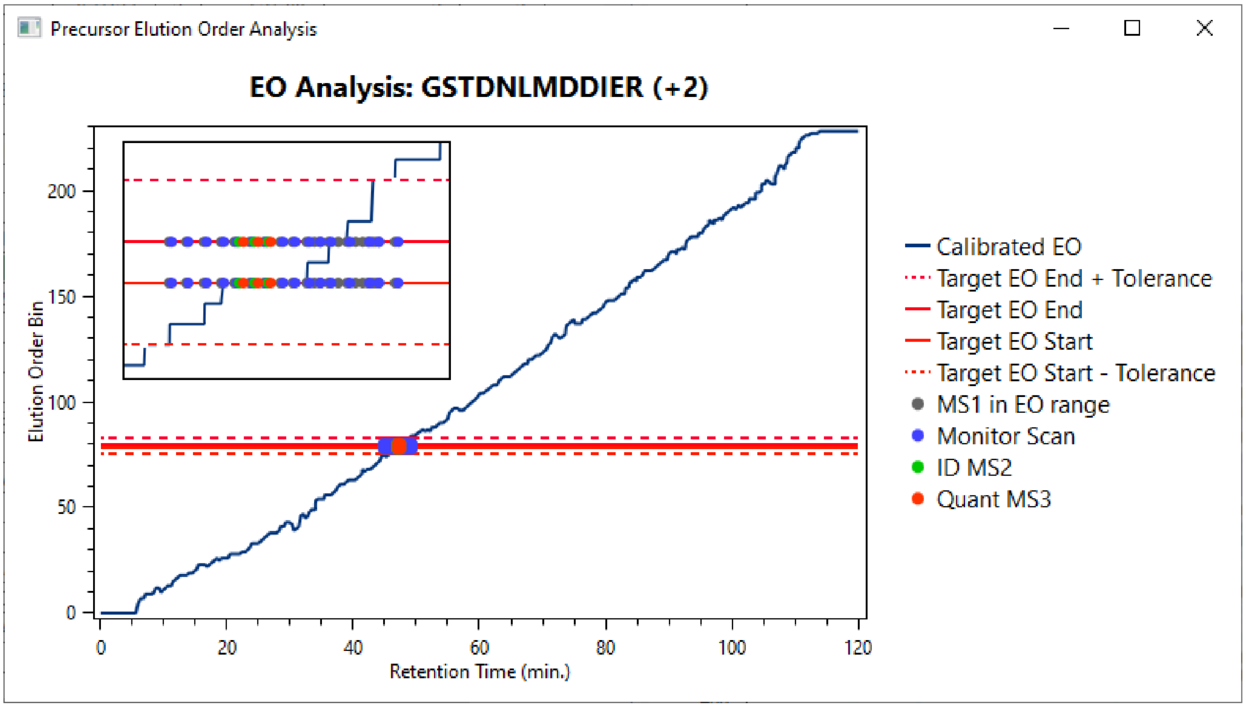


**Figure S6. The Target EO Analysis pane in GoDig Viewer.** The same plot as Figure S5 with the following exceptions: only scans pertaining to the selected target are plotted; MS1 scans acquired while within EO range, monitor scans, and unsuccessful IDMS2 scans are additionally plotted; and the library EO bin range of the target as well as the EO range established by the tolerance parameter are illustrated as horizontal red lines. Inset: a zoomed-in region of the same plot.

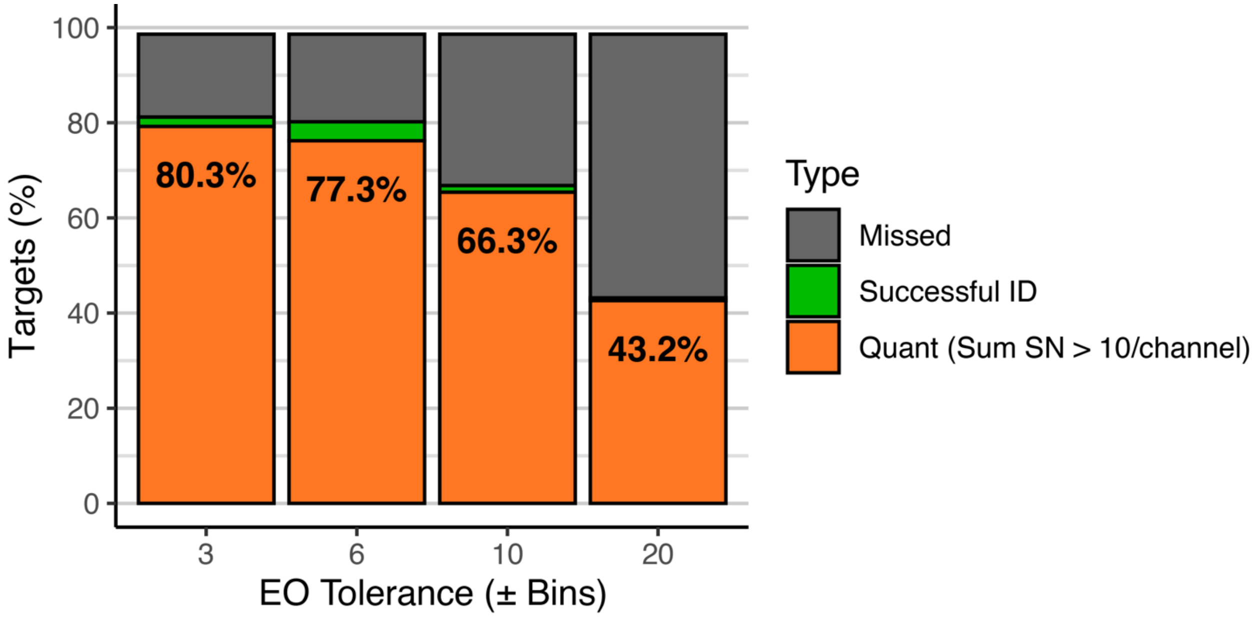


**Figure S7. Effect of elution order (EO) Tolerance on success rates.** Though increasing EO tolerance can enable detection of some out-of-bounds targets, the total success rate can decrease, as shown by this analysis. One bin corresponds to 0.5 min on average. Data are from single-injection GoDig analyses targeting 500 “super fliers” (see main text) with different EO tolerances.

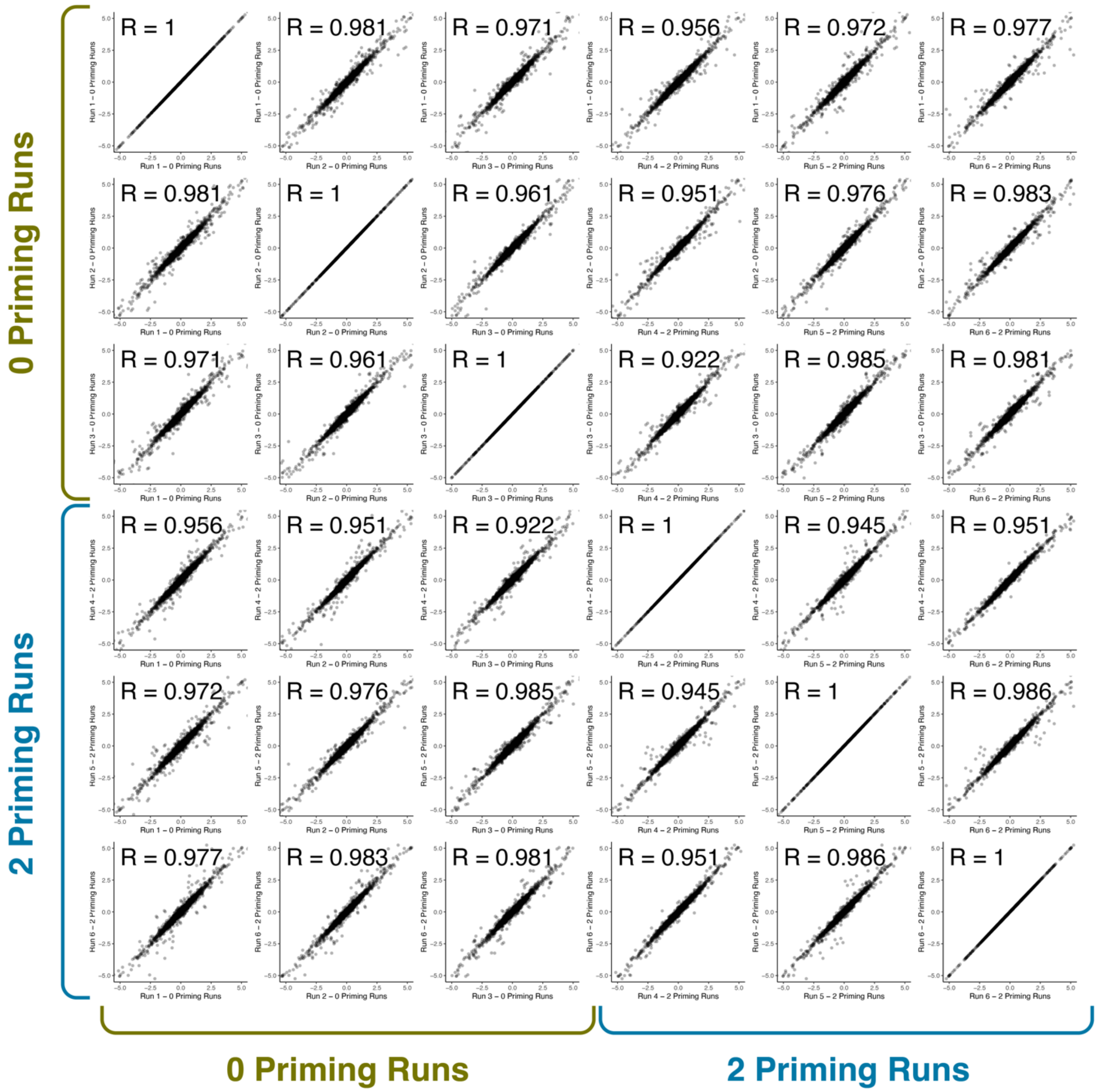


**Figure S8. Scatter plots comparing peptide quantity fold changes between cell lines.** Each scatter plot is a comparison between two analytical runs from Figure 2D. Each point represents a log-transformed fold change for a peptide between two different cell lines. The $\log_2$(fold change) from one run is plotted along the x-axis and the same $\log_2$(fold change) from another run is plotted along the y-axis. In the left half of the grid, no priming runs were used for measurements along the x-axes, whereas in the right half, two priming runs were used for the x-axes and all targets were primed. In the top half of the grid, no priming runs were used for measurements along the y-axes, whereas in the bottom half, two priming runs were used for the y-axes and all targets were primed. Each R is the Pearson correlation coefficient between the two runs.

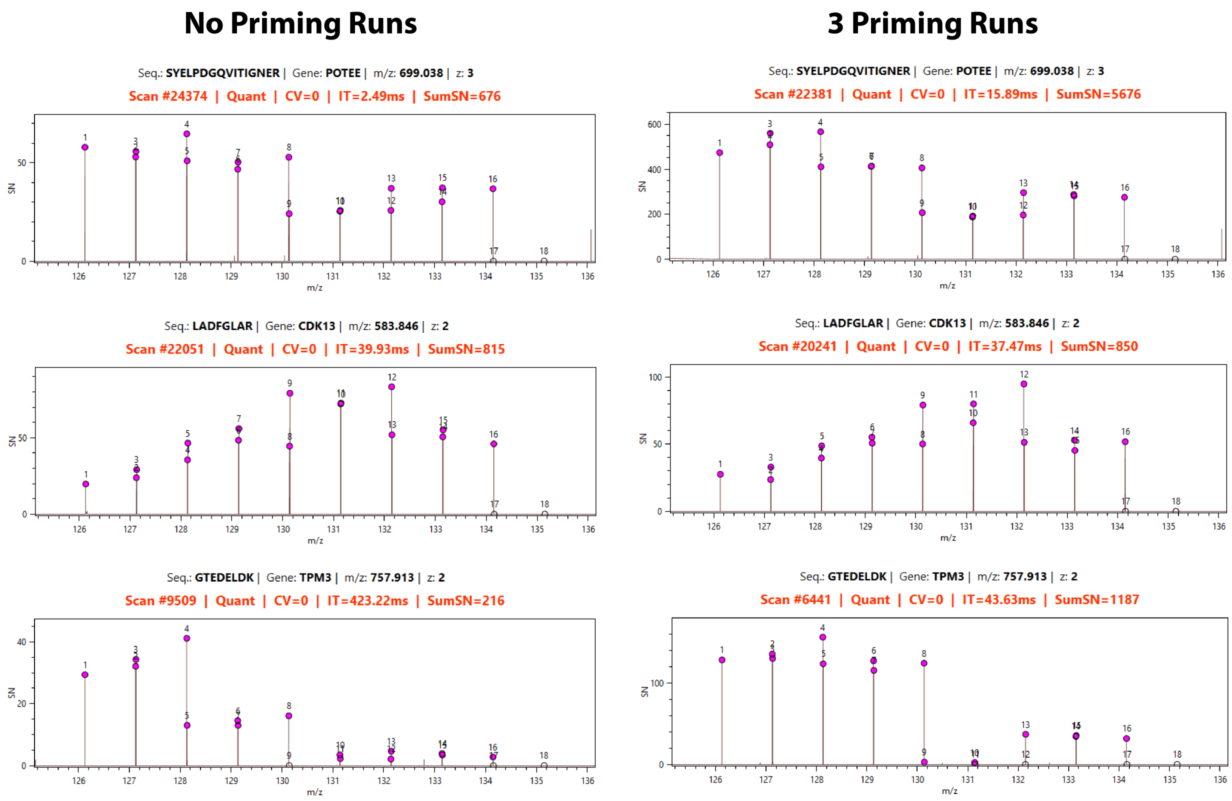


**Figure S9. Selection of MS3 spectra acquired with and without priming runs.** Each peak is labeled with the TMT channel index. Y-axis is intensity divided by orbitrap noise. Channels 17 and 18 are zero because the corresponding reagents were not used in the experiment.

| Run | IDs | Quants | IDs (%) | Quants (%) |
|---|---|---|---|---|
| 1 | 399 | 394 | 99.8% | 98.5% |
| 2 | 398 | 395 | 99.5% | 98.8% |
| 3 | 396 | 390 | 99.0% | 97.5% |
| 4 | 392 | 387 | 98.0% | 96.8% |
| **Mean** | **396.3** | **391.5** | **99.1%** | **97.9%** |
| **SD** | **3.1** | **3.7** | **0.8%** | **0.9%** |

**Figure S10. GoDig results targeting 400 super flyers in 250 $\mu$g of 4-cell-line sample for 2 h with 2 priming runs and 4 consecutive analytical runs.** "Quants" = targets quantified (MS3 triggered and TMT sum signal-to-noise > 160).

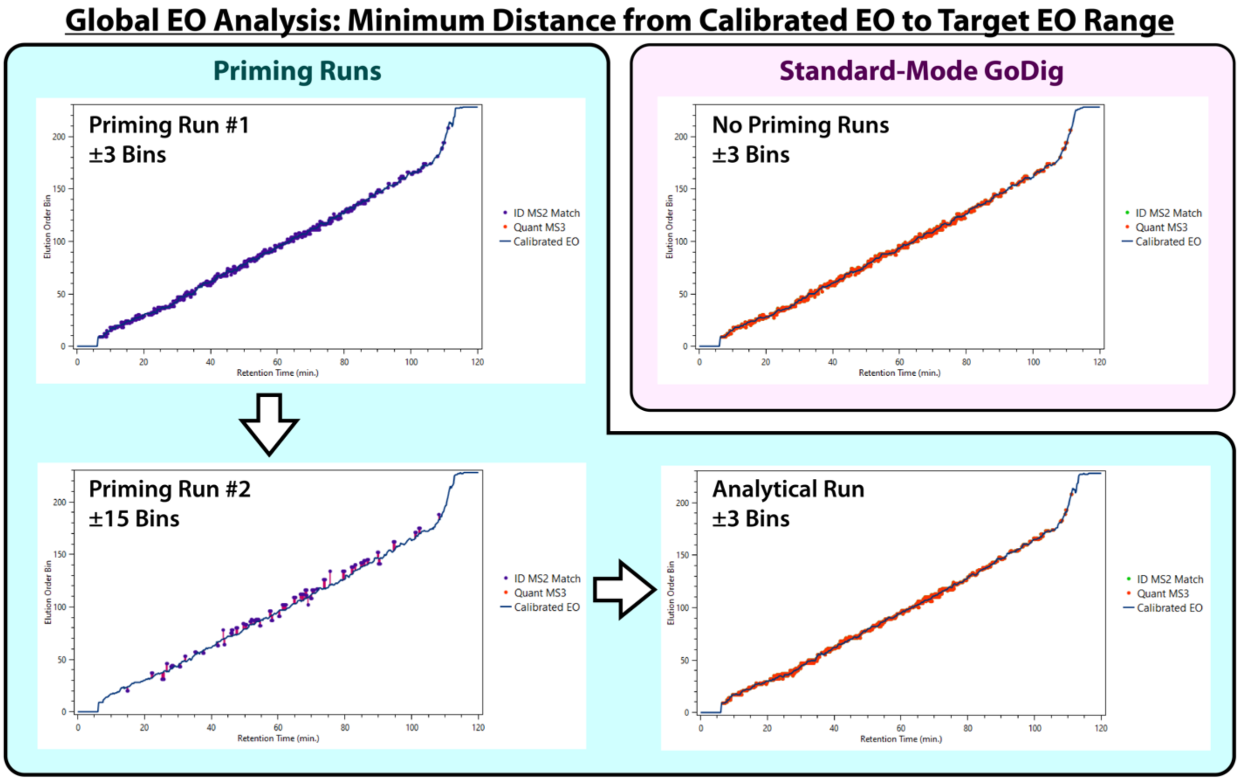


**Figure S11. Same as Figure 4 except the y-axis plots the EO bin in each library target EO range that is closest to the current calibrated EO bin.** I.e., if the current calibrated EO bin lies within the library target EO range, the current calibrated EO bin itself is used, and if not, then the closer of the two EO bins defining the range is used.

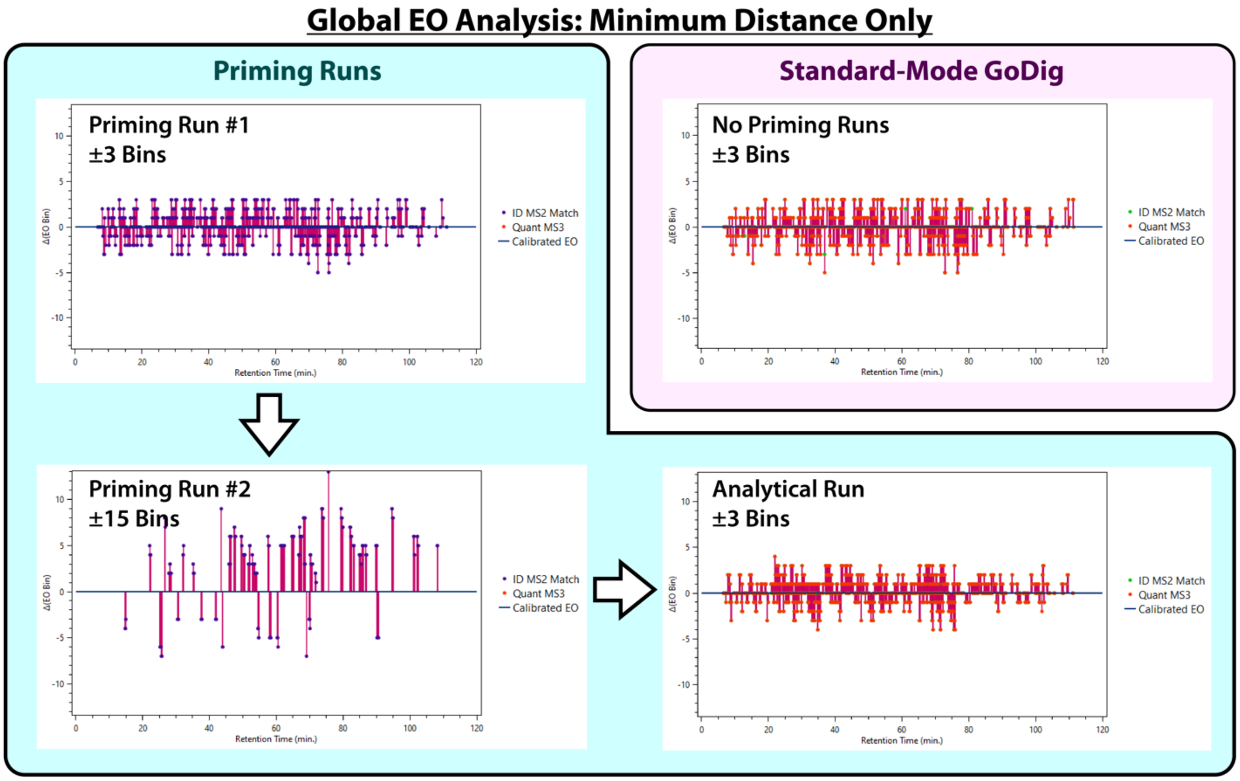


**Figure S12. Same as Figure S11 except the y-axis plots the difference between the library target EO bin and the current calibrated EO bin.**

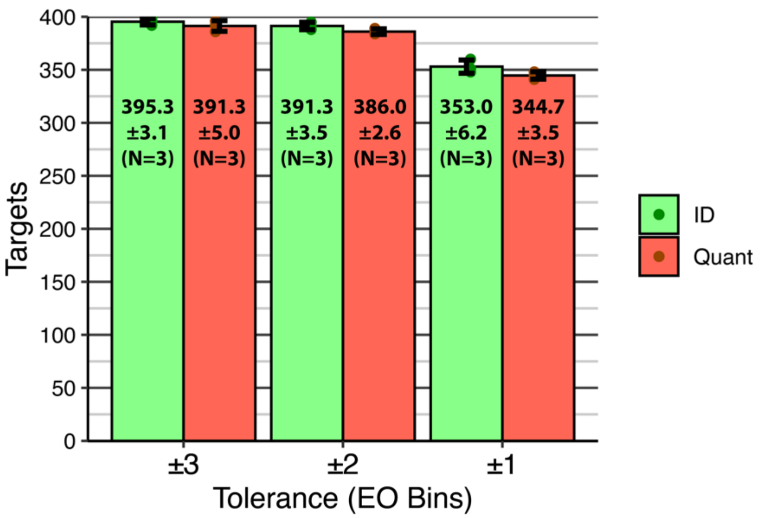


**Figure S13. Priming runs enable narrower EO windows for analytical runs.** The ±3-bin data were acquired in the same way as Figure 2D except that a single set of 2 priming runs was performed and followed by 3 consecutive analytical runs, all using the same corrected EO bins. The ±2-bin data and ±1-bin data are each a set of 3 analytical runs using the same corrected EO bins exported by GoDig during the ±3-bin experiment shown here.

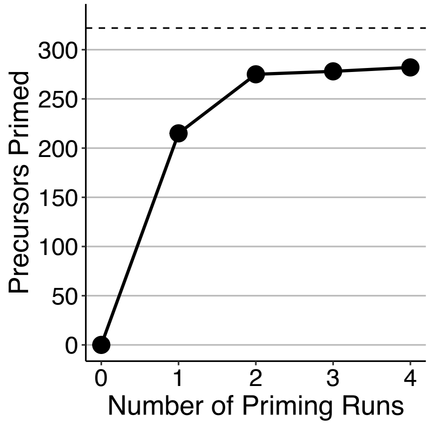


**Figure S14. Priming runs targeting 322 precursors matching 161 macroautophagy genes.** Over 4 priming runs, a total of 282 precursors were primed. Within these, 125 pairs of precursors matching the same genes were primed (125 genes = 250 precursors).

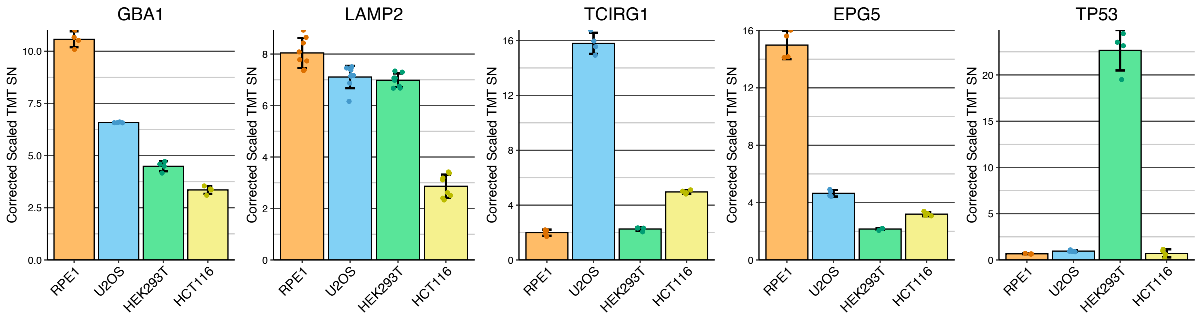


**Figure S15. Reanalysis of GoDig data from Yu et al. 2023 (Ref. 1).** As described in Ref. 1, the same 4-cell-line mixture as described in this work was prepared and then fractionated using HPLC at basic pH. The fractions were analyzed using an untargeted RTS-SPS-MS3 method to generate these data.
